# Convolutional Neural Networks and Outline Analyses for Archaeobotanical Studies of Domestication and Subspecific Identification

**DOI:** 10.1101/2023.09.15.557939

**Authors:** Vincent Bonhomme, Laurent Bouby, Julien Claude, Camille Dham, Muriel Gros-Balthazard, Sarah Ivorra, Angèle Jeanty, Clémence Pagnoux, Thierry Pastor, Jean-Frédéric Terral, Allowen Evin

## Abstract

The identification of archaeological fruits and seeds is crucial for understanding the relationships between humans and plants within the cultural and biological history of both wild and cultivated species. We compared the relative performance of a deep learning approach, namely convolutional neural networks (CNN), and outline analyses via geometric morphometrics using elliptical Fourier transforms (EFT) at identifying pairs of plant taxa. We used their seeds and fruit stones that are the most abundant organs in archaeobotanical assemblages, and whose morphological identification, chiefly between wild and domesticated types, allows to document their domestication and biogeographical history. We used existing modern datasets of four plant taxa (barley, olive, date palm and grapevine) corresponding to photographs of two orthogonal views of their seeds that were analysed separately to offer a larger spectrum of shape diversity. Sample sizes ranged from 473 to 1,769 seeds per class, which constitute a relatively small dataset for training CNNs models yet typical within archaeobotanical research. On these eight datasets, we compared the performance of CNN and EFT coupled with linear discriminant analyses. Our objectives were twofold: i) to test whether CNN can beat geometric morphometrics in taxonomic identification and if so, ii) to test which minimal sample size is required. We ran simulations on the full datasets and also on subsets, starting from 50 images in each binary class. For the CNN network, we deliberately used a candid approach relying on pre-parameterised VGG19 network. For EFT, we used a state-of-the art morphometrical pipeline. The main difference rests in the data used by each model: our CNN used bare photographs where EFT used outline coordinates. This “pre-distilled” geometrical description of seed outlines is often the most time-consuming part of morphometric studies. Results show that our CNN beats EFT in most cases, even for very small datasets. We finally discuss the potential of CNNs for archaeobotany, and how bioarchaeological studies could embrace both approaches, used in a complementary way, to better assess and understand the past history of species.

## Introduction

From Aristotle to Darwin, the form of organisms has long inspired our understanding of the living world. In some disciplines such as archaeobotany, the shape of plant remains is, most often, the only available datum. Both qualitative and quantitative morphological criteria first allowed to identify plant remains, particularly seeds and fruit stones, often at the species level (Jacomet, 2008; Zohary, Hopf, & Weiss, 2012). Then, purely quantitative tools, and chiefly geometric morphometrics, allowed for finer-grained, statistically assessed identifications, to further explore the morphological size and shape variation.

Geometric modern morphometrics (further abbreviated GMM), is the statistical description of shape and its covariation (Kendall, 1989). It uses generic mathematical transformations to convert shape and size into quantitative variables. Most GMM studies either uses configuration of landmark coordinates, the geometry of curves (closed or not) or, more recently, entire surfaces. Curves analyses are often favoured in archaeobotany due to the absence of clear landmarks, if any, on botanical organs and elliptical Fourier transforms (further abbreviated EFT) is the most popular approach.

By comparing archaeological material to modern collections of reference, GMM and EFT in particular for plants, allowed fine-grained inferences, in particular to document the emergence of new morphological types, evidence domestication syndromes, reconstruct the dynamics of their diffusion in both time and space, and overall gain insights into the intertwined histories of human societies and domesticated plants (Bonhomme et al., 2017a; Bonhomme, Terral, et al., 2021; Bouby et al., 2013; Bourgeon et al., 2018; Burger, Terral, Ruas, Ivorra, & Picq, 2011; Evin et al., 2022; Jesus et al., 2021; Pagnoux et al., 2015; Ros, Evin, Bouby, & Ruas, 2014; Roushannafas, Bogaard, & Charles, 2022; Tarongi et al., 2021; Terral et al., 2004, 2012, 2010; Wallace et al., 2018).

Deep learning quickly became a game-changer from academia to industry, through its versatility and cutting-edge achievements. Computer vision in general has largely benefited the synergy between the massive democratization of computational power and the arrival of software frameworks on top of solid mathematical foundations. Convolutional neural networks (further abbreviated CNN, (Lecun, Bottou, Bengio, & Haffner, 1998), in particular, have been at heart of very diverse supervised classification tasks, from autonomous vehicles to plant identification (Alzubaidi et al., 2021; Berganzo-Besga, Orengo, Lumbreras, Aliende, & Ramsey, 2022). However CNN still remain relatively rare in paleontological and archaeological studies (Soroush *et al*., 2020; Romero *et al*., 2020; Garcia-Molsosa *et al*., 2021; Loddo *et al*., 2021; and the review by Bellat *et al*., 2025) and also in morphometrics (Le, Beurton-Aimar, Zemmari, Marie, & Parisey, 2020; Miele, Dussert, Cucchi, & Renaud, 2020) yet datasets of large number of images are now available and can be employed to develop new tools for specialized tasks like seed recognition (Yuan et al., 2024).

Date palm (*Phoenix dactylifiera* L.), grapevine (*Vitis vinifera* L.) archaeological seeds, barley (*Hordeum vulgare* L.) caryopsis and olive (*Olea europaea* L.) stones have been intensively studied in archaeobotany using geometric morphometrics. They are four important taxa of human subsistence in the Mediterranean basin since millennia. The presence of the wild progenitors of the domestic forms in vast geographic ranges makes the identification of the wild or domestic status of the archaeobotanical remains of date palm, olive and grapevine particularly difficult. In addition, the presence of multiple types for barley in the region, exploited for diverse use and with different agricultural practices require intra-specific identification.

The morphological distinction between wild and domestic types using GMM is now very accurate for olive (Terral et al., 2004, 2021) and grapevine (Bonhomme et al., 2022; Terral et al., 2010). On the other hand, distinguishing between wild and domestic date palm seeds (Gros-Balthazard et al., 2017; Terral et al., 2012), as well as between two- and six-row barley grains remains challenging (Bonhomme et al., 2017b; Jeanty et al., 2024; Ros et al., 2014; Wallace et al., 2018).

In that context, this paper aims to test the potential of a deep learning approach for archaeobotanical identification and ask the following questions: i) can a CNN outperforms baselines obtained with GMM and if so, ii) how much data are typically required to train the models? Here, we used four plant models presenting binary challenges below the species level, at core in archaeobotanical studies. More precisely, our aim was to distinguish between wild and domesticated types of date palm, olive and grapevine, and between two- and six-row barley,

A CNN model correctly trained on large datasets is expected to outperforms EFT approaches, providing taxonomical differences are reflected in some morphological contrasts, at least because EFT are limited to the geometrical differences of outlines, while CNN applied on images can capture any morphological discriminant feature beyond shape, texture for example. That being said, several conditions of our models make such expectation far from granted here:

i. *Low inter-class differences*: differences tested here, chiefly shape differences, ranged from subtle at best to extremely challenging; the group labelling was certain only because the identification was obtained through molecular markers (for the date palm) or on entire plants cultivated in biological conservation centres (other models).
ii. *Small sample sizes*: the available datasets were particularly small compared to those usually deployed in CNN learning tasks. The datasets used here were obtained through 2D images acquired following rigorous and time-consuming protocols, which limit the number of biological objects that can be analysed in the context of archaeobotanical studies.
iii. *Challenging baselines*: existing baselines obtained through GMM are already good to very good.
iv. *Accessible models*: our intention was to develop CNN-based pipelines, reasonably easy to run by non-expert users using general-purpose computers.
v. *Taphonomic biais*. Charring, desiccation or waterlogging related to the fossilisation of the fruit and seed stones can potentially generate important sampling bias in CNN by comparison to GMM as the first one might focus on texture rather than outline geometry.

We first present the models used and compare their performance to geometric morphometrics. Finally, we discuss the pros and cons of CNN versus EFT and propose an agenda of future researches.

## Material and methods

### Statistical environment

All analyses were run using R 4.1.3 (R Development Core Team, 2024). We used a MacBook Pro 2013 model with a 2,6 GHz Intel Core i5 CPU and 16 Go 1600 MHz DDR3 RAM. Data manipulation and visualization was done using *tidyverse* 2.0.0 (Wickham et al., 2019). Image manipulation was done using *magick* 2.7.3 (Ooms, 2021). All morphometric analyses were performed using *Momocs* 1.4.1 (Bonhomme, Picq, & Claude, 2020; Bonhomme, Picq, Gaucherel, & Claude, 2014) and linear discriminant analyses using *MASS* (Venables & Ripley, 2002). CNN models used *keras* 2.15.0 (Allaire & Chollet, 2020), the R interface to the eponym Python 3.7 architecture, which here ran on CPU alone.

### Datasets used

Among the model species studied by our team, we retained those for which we have enough material, secure identification and associated publication record: grapevine pips (Bonhomme, Picq, Ivorra, et al., 2020a; Pagnoux et al., 2015), barley grains (Jeanty et al., 2023), olive stones (Bourgeon et al., 2018; Terral et al., 2021) and dates seeds (Gros-Balthazard et al., 2017; Terral et al., 2012) (Table 1). All models corresponded to a binary classification task with 2-versus 6-row types of barley (*Hordeum vulgare*), and wild versus domestic for the three other taxa. These datasets only comprised modern material from collections of references, commonly used to compare with archaeological material.

**Table 1:**
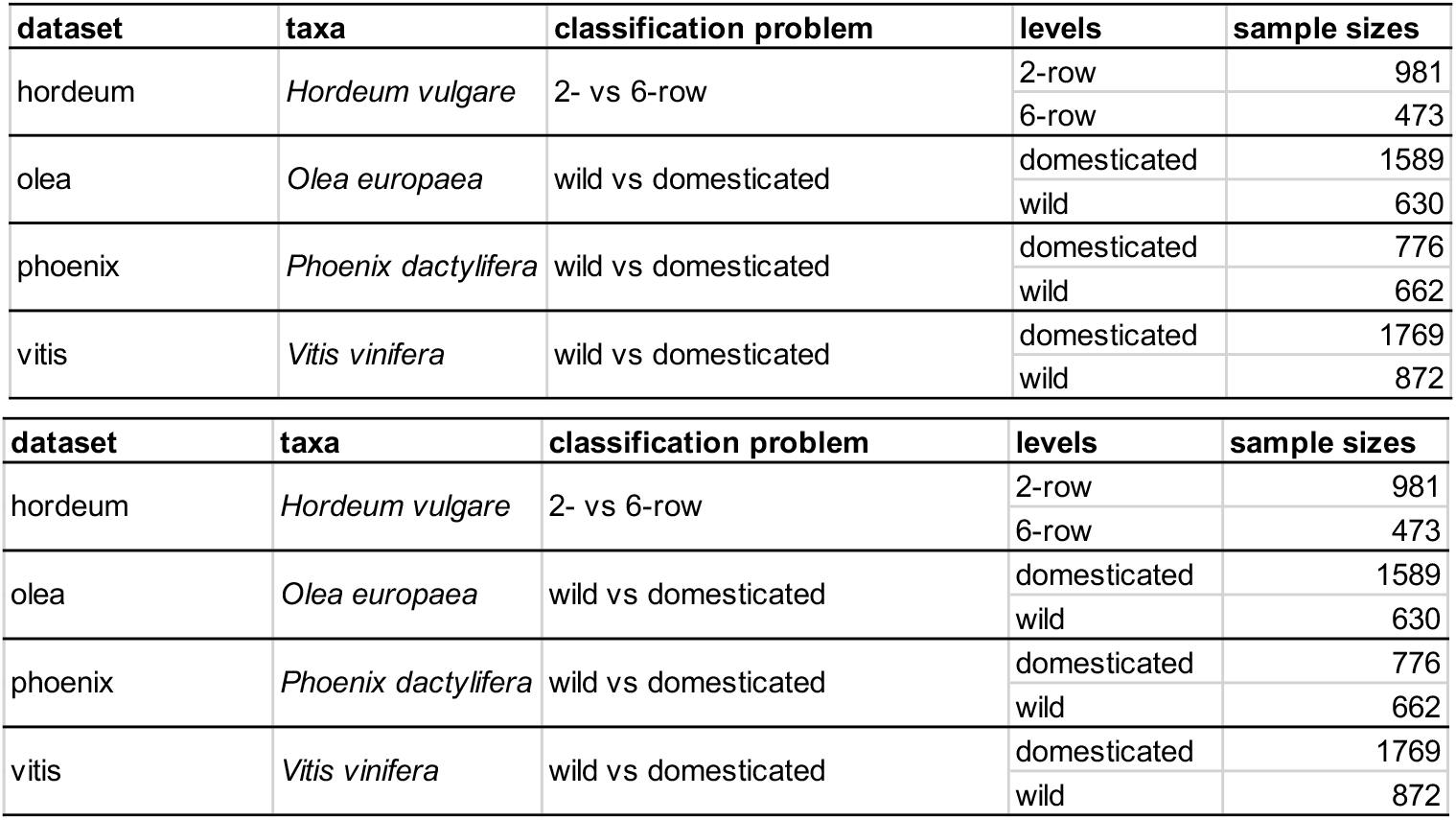
Material used. Each of the four taxa provided two views treated separately.

All seeds/stones/fruits were photographed in dorsal and lateral views using a stereomicroscope coupled with a digital camera. It is worth noting that GMM identification is usually obtained by combining the information brought by the two orthogonal views but we chose here to *not* combine these views to increase the number of “independent” datasets and have a larger spectrum of shapes (Figure 1).

**Figure 1:**
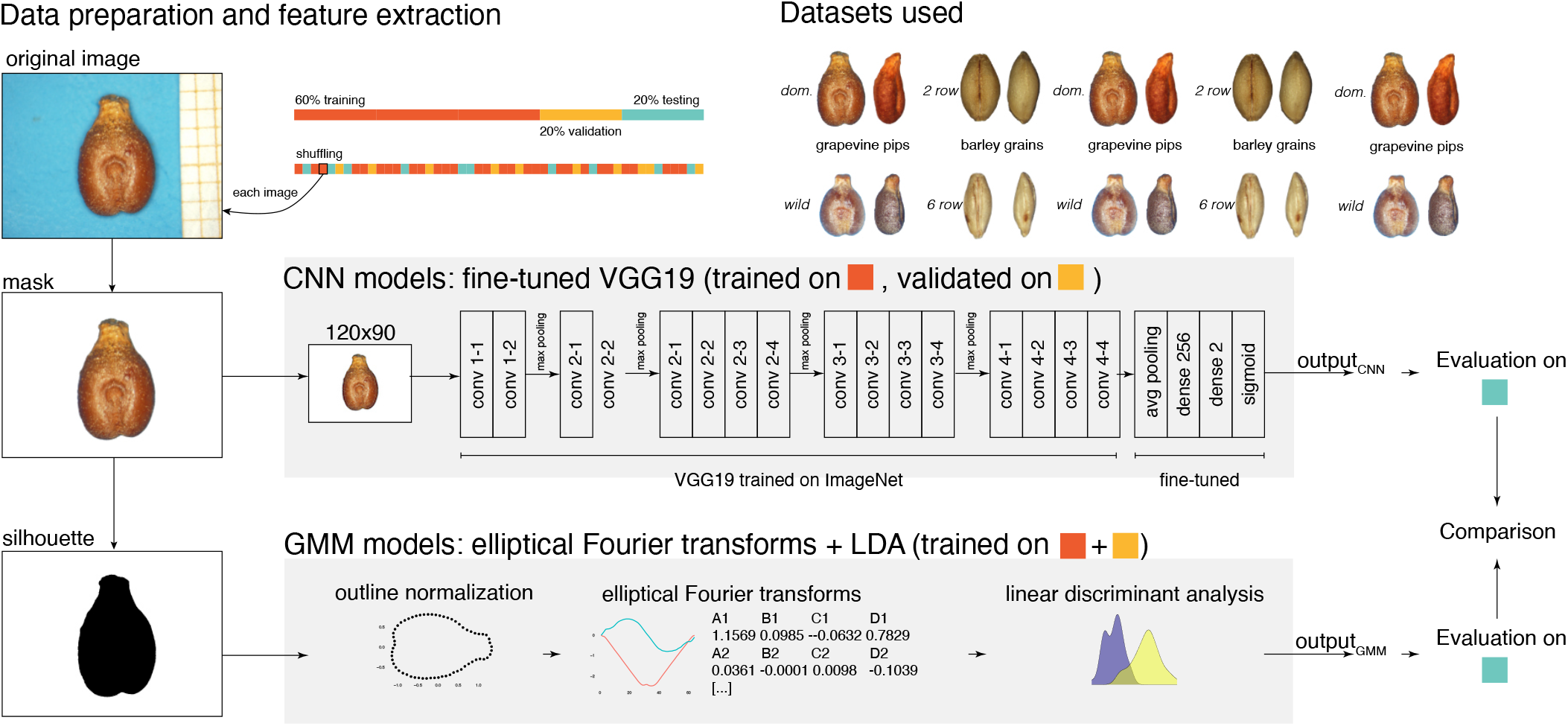
Schematic description of the GMM and CNN models used. For each taxon, an archetypical individual seed is presented.

### Convolutional neural networks

Our CNN models used the VGG19 architecture (Simonyan & Zisserman, 2014) with the weights trained on the *ImageNet* reference dataset (Deng et al., 2009), as available in *keras*. The convolutional base, with feature hierarchies learnt on *ImageNet*, was frozen. Given we did not want to predict *ImageNet* classes, the last three dense layers were replaced with two fully connected dense layers and only these last layers were fine-tuned. The first has 32 units and a rectified linear unit (ReLU). Because all models were binary classification, the last layer has two units and a sigmoid activation (Figure 1). The loss was calculated using binary cross-entropy for binary classification tasks. We used two callbacks to control the training step. The first controlled the learning rate, based on loss decrease, initially fixed to 10^-2^ with a decay factor of 10, a patience of 10 epochs and a minimal value of 10^-7^. The second stopped the training with a patience of 20 epochs with no accuracy improvement. These two callbacks were used to homogenise training among models.

For each dataset, the number of images was balanced between classes, using a random sampling without replacement among available images (Table 1). This allowed to explore the effect of different sample sizes on final CNNs performance. For each sample size tested, 60% of the total number of images was used for the training set, 20% for the validation set and the last 20% for evaluating the model. The training set was used to adjust weights while the validation set was used to evaluate model performance back-propagate results to the unfrozen layers at the end of each epoch. The evaluating set was used only once and after the training step, to report the model performance on images never seen before by the model.

The rgb images were converted to grayscale with pixel values standardized between 0 and 1 and reduced to a resolution of 120×90 pixels, while maintaining the aspect ratio (Figure 1). The first layer of the VGG19 convolutional based was adapted accordingly.

### Geometric morphometrics baseline

We used outline analysis using elliptical Fourier transforms (EFT) (Bonhomme et al., 2014; Claude, 2008; Kuhl & Giardina, 1982). We first converted full-sized images into silhouette masks on which 360 outline coordinates were sampled, equally spaced along the curvilinear abscissa. We then normalized outlines for their size, position, rotation and first point and obtained enough harmonics to gather 95% of the total harmonic power (6 for all datasets). Then, a linear discriminant analysis (LDA) was trained on the same dataset as for CNN yet combining the training and validation sets (Figure 1). The general methodology is detailed elsewhere (Bonhomme et al., 2014; Bonhomme, Picq, Ivorra, et al., 2020b).

### Model comparisons

Ten replicates were used for each of the eight datasets and each was tested with increasing sample sizes (Table 1 and Table 2). Given one of the 560 runs, the very same sets of images (or masks) was submitted to both CNN and EFT. The only difference is that, for EFT, the two training and validation sets were combined for training then evaluated on the same 20% as for CNN (Figure 1). This cross-validation scheme allowed direct comparisons between the respective performances of each model. Performance was measured with accuracy, that is the proportion of correctly identified individuals. Sensitivity and specificity were also calculated.

**Table 2:**
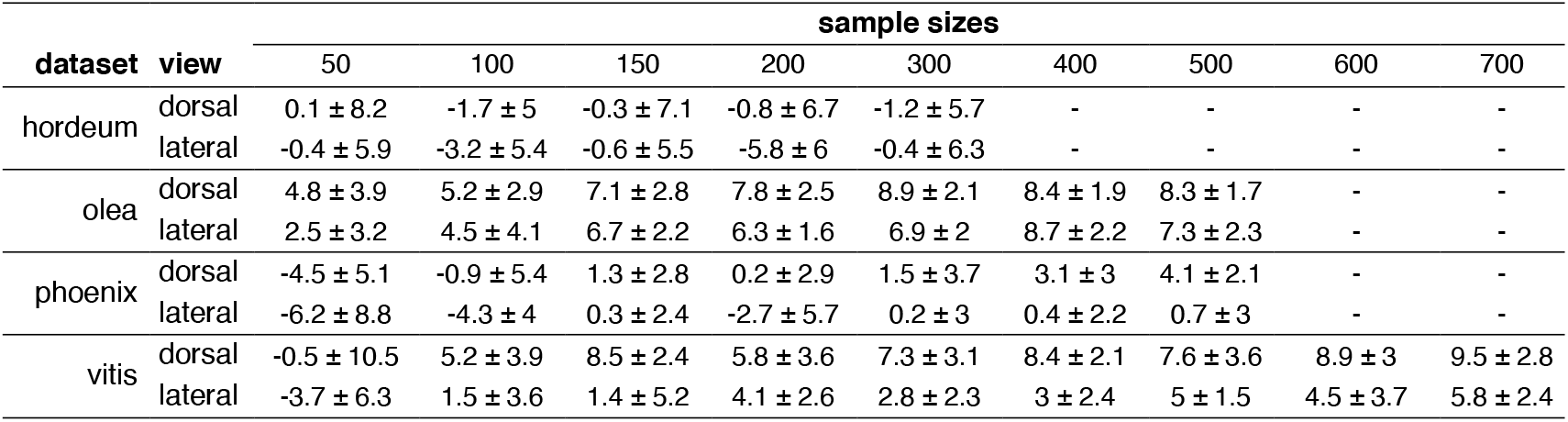
Mean accuracy (CNN – EFT) differences ± standard deviation, expressed as percentages, for each of the 10 replicates. Sample sizes are expressed as the total number of images used per class (training/validation + evaluation). Cells with ‘-’ could not be calculated due to sample size limitations.

## Results

In most cases, CNN beat EFT (419 cases over 560, that is 75% - Figure 2, Figure 3, Table 2). This is particularly true for larger training sets.

**Figure 2:**
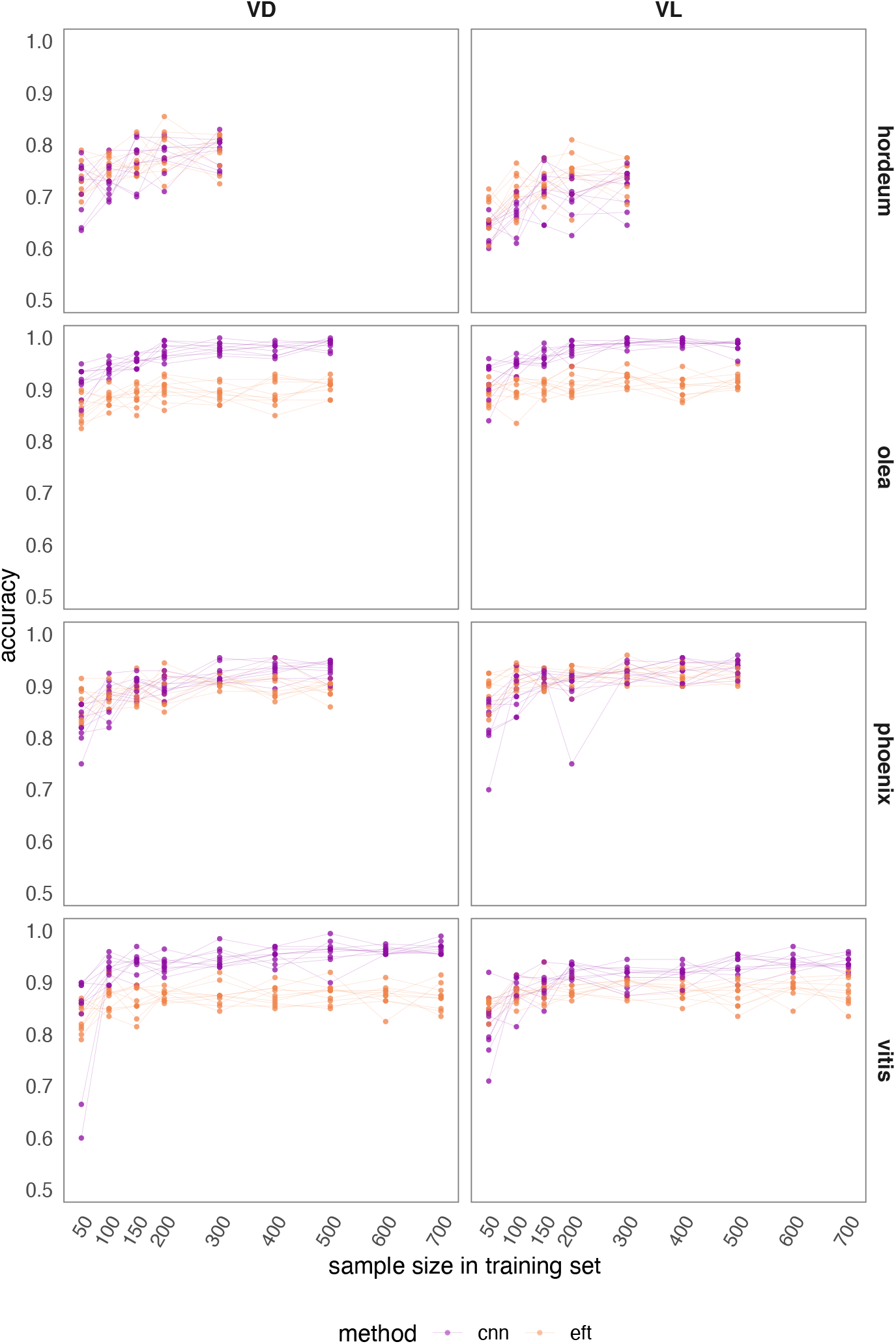
Model performances for each dataset presented using accuracy, training sample sizes and replicates for CNN and EFT. Sample sizes are expressed as the total number of images used per class (training/validation + evaluation). The models are run for the ventral (VD) and lateral (VL) views. Two-row vs six-row barley (hordeum) are compared, as well as the wild and domestic forms of olive (olea), date palm (phoenix) and grapevine (vitis). The same graphs for precision and recall are available in the supplementary material and show similar trends.

**Figure 3:**
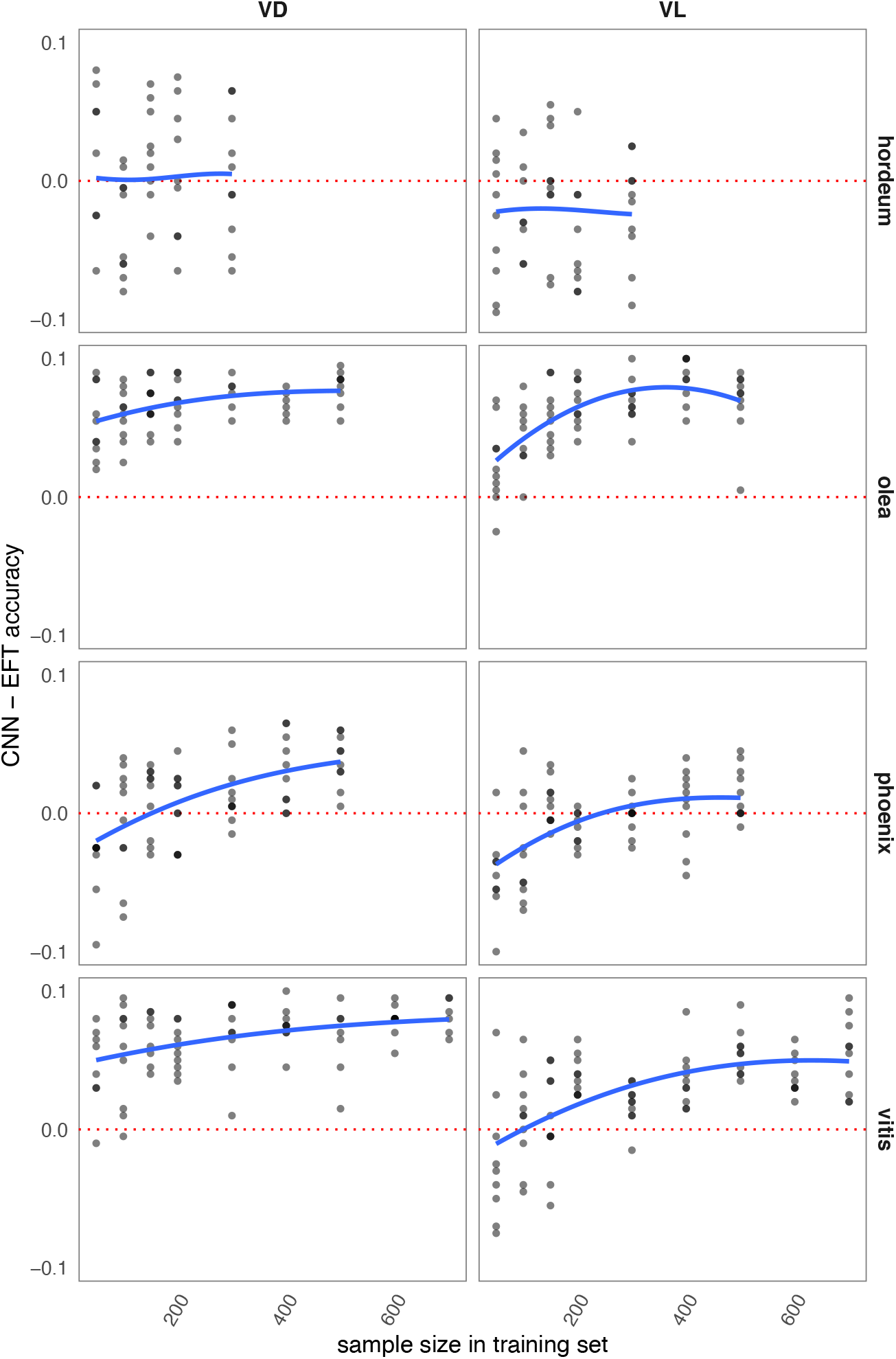
Model performances for each dataset training sample sizes and replicates, presented as absolute CNN - EFT accuracies. Sample sizes are expressed as the total number of images used per class (training/validation + evaluation). The models are run for the ventral (VD) and lateral (VL) views. Two-row vs six-row barley (hordeum) are compared, as well as the wild and domestic forms of olive (olea), date palm (phoenix) and grapevine (vitis).

Among the eight datasets (further referred using their vernacular name) two groups can be made: olive and grapevine in one hand, barley and date palm in the other, no matter the view considered. For grapevine and olive, where EFT “already” provided good accuracies, CNN perform even better, particularly for the large sample sizes. For grapevine with 700 images, average CNN accuracies range from 94 to 99% for dorsal view and from 92 to 96% for lateral view. For olive with 500 images, performances range from 98 to 100% for dorsal view, and from 97 to 99% for lateral view. For barley and date palm, the results seem more mitigated at first glance (Figure 2), yet, on average (Table 2) the CNN also achieve better accuracies when the datasets are large enough. For sample sizes above 150 individuals, CNNs are better in most cases for barley and consistently for date palm. These two groups of results are reflected in the mean differences between models for the largest sample size: olive and grapevine gained ∼10% accuracy where date palm and barley gained less than 5%. The patterns observed in sensitivity and specificity mirrored those of accuracy and are available in the ESM. Finally, to give an idea of computational time, a single iteration of the 56 models pairs took ∼17 hours to complete, with less than 1% dedicated to the EFT. In the other hand, post-treatment time for preparing pictures is virtually zero for CNN and about 1 min per picture for EFT, that is about a full-time week for each dataset here.

## Discussion

Our results show that even a candid CNN approach could outperform state-of-the-art EFT to identify plant seeds and fruits below the species level. Even if the performance boost is not dramatical for all four studied taxa, this was a quite surprising result since the CNN beat almost consistently our EFT baselines even when the sample sizes were small.

Regarding the four pairs of taxa studied, identifying wild and domestic types for olive and grapevine is relatively easy using the seed shape but distinguishing between the wild and domestic date palm, and the two- and six-row barley is challenging, not to say troublesome. For hordeum (lateral view), CNN are even, on average, below the EFT obtained with accuracy. Further research using refined CNNs architectures would be helpful on that particular dataset.

Here, when geometrical differences between studied pairs are quite obvious macroscopically (olive and grapevine), the CNN clearly beat GMM identification and is close to perfect when the sample sizes of the training sets exceeded 500 hundred seeds. For instance, over the 10 replicates, a single olive stone in lateral view was wrongly identified among the 1400 evaluated images (20% of 10*700). Using outline analyses, accuracies around 95% can now be reached for certain taxa (e.g. olive and grapevine), particularly when combining several views (Bonhomme, Picq, Ivorra, et al., 2020b), but here CNN only have raw 120×90 images as inputs.

Perhaps the most surprising result is that CNN also beat EFT baseline even when trained with only ∼100 images in each class, at least for these two “easy” models (here grapevine and olive). Given how costly and time-consuming is the constitution of a reference collection, this means that CNN can be tested early and possibly cut off these costs. Also, that methods applied here could be easily tested in many other archeological models whether they are plant, animal organs or non-biological artefacts, imprints, etc.

One important result here is that CNN can still improve their classification score when increasing the training sample size well after the classification score of GMM can no more be improved (because of limitations associated with the linear discriminant analyses or because the number of available variables is limited). Our results seem also to indicate that GMM and linear discriminant analysis allow to fast reach their maximal accuracy but rapidly reach a plateau (corresponding to 50 to 150 analyzed individuals). With larger sample size, they are clearly performing less well than CNNs.

Deep learning approaches are now quite common for animal and plant species identification, particularly for citizen science projects (Picek, Šulc, Patel, & Matas, 2022; Willi et al., 2019), but remain so far very new when it comes to archaeological material (but see Miele *et al*., 2020) or morphometrics (but see Le et al., 2020). To the best of our knowledge, this is the first time CNNs are used for such sub-specific identification task in plants, a fortiori on four different model taxa. The results shown here appeal to further studies to test how they could be extended to other archaeological material, other plant or animal taxa and at the species level. Here we show that, at least in some cases, the diversity at even lower taxonomic levels can be explored. This would be of prime interest to develop tools that can be used not only by archaeobotanists but also by any people interested in identifying variety (*e*.*g*. for conservation purposes). In palynology, another field that may be developed in an archaeological context, deep learning using CNNs has already proven to be helpful in the fastidious task of pollen and phytolith identification and counting (Berganzo-Besga et al., 2022; Gimenez et al., 2024; Sevillano, Holt, & Aznarte, 2020).

In this paper, our main intention was to take the archaeobotanists point of view: “How can my reference collection help me interpret the identity and significance of my unknown seeds?”. Despite these encouraging initial results, it remains important to note the most apparent potential pitfalls.

First, CNN and EFT as implemented here neither work with the same information (CNN use images, EFT use an outline geometry), nor use the same method here for the classification (CNN use a sigmoid activation, EFT uses LDA). EFT is, by construction, limited to the description of the shape and form variations where CNN use a number of other variables that can be useful (and also possibly detrimental) for classification, such as colour, texture, or patterns that go beyond the outline geometry. This calls for additional research, for the sake of a more direct comparison between paths. For example: how would CNNs (and other deep learning tools) behave with more or less “distilled” geometric information when given raw images, cropped images, masked images, (x, y) outline coordinates, EFT coefficients or even PC scores? Also, would an intermediate segmentation model mask (i.e. with texture) seeds be interesting in terms of robustness and performance? Future research will help clarify how CNN, and more largely deep learning can really be a game changer in archaeobotanical studies. Second, CNN models used here may be highly sensible to different image acquisition environment, including apparatus, lightning, operator, post-processing, etc. Such biases have already been investigated, sometimes with workarounds (Da Rin et al., 2022; Fortin et al., 2018; Kothari et al., 2014). Datasets used here were obtained in multiple settings and environments in the last twenty years and further experiments will likely share the same potential pitfall, either directly by combining cross-laboratory acquisition, by using the reference datasets provided with this paper, or by evaluating archaeological material with models trained on modern material. This may also call for other approaches using masks or outlines which are often already obtained for further outline analyses anyway. Additional work to test for these potential biases is needed.

More generally, should we expect rivalry or synergy emerge between CNN and geometric morphometrics? For rough identification, CNN will likely become a more popular tool for future archaeobotanical studies, and possibly the next standard toolkit. Here, we insist on the fact that our CNN architecture was deliberately kept simple for both practical and conservative reasons: we had many models to run that needed to be generic and the point was to test if a candid CNN approach could beat state-of-the-art EFT. There is definitely room for improvement by using better models, fine-tuning them, larger datasets, larger images and by combining views or even using 3D models of the objects. That being said, with the best will in the world a model cannot see what simply does not exist. In some cases, a single ratio of lengths can achieve nearly perfect identification when morphological differences are trivial. This is the case for example for grapevine (Bonhomme et al., 2022). On the other hand, meaningful differences for human use may just not be reflected on the studied organs. Somewhere between these two extremes are a wide range of real differences that can only be detected by statistical means (Bonhomme, Ivorra, et al., 2021; Bonhomme, Terral, et al., 2021). This is where methodological refinement makes the most sense and a natural playground for deep learning approaches.

EFT have the advantage of translating the shape into coefficients that can be directly treated as quantitative variables. Also, the inverse transformations are mathematically defined, so that one can go “back” to shape from coefficients, which allows rich insights into the morphological space of taxa of interest and the comparison between the relative occupancies between taxonomic, diachronic or synchronic assemblages. For that matter, the best equivalent CNN have to offer so far are activation maps where one can visualize for each image, the regions that triggered the final vote. Even though the reputation of being black boxes is largely erroneous, CNN are and will likely remain less handy to that respect mostly because they use images that are difficult to interpret in terms of model explainaibility than, say, outline coordinates. More generally, “to predict is not to explain” (Thom, 2010), and in our opinion, CNN and EFT should be seen as complementary approaches rather than competitors. Future studies will explore this assumption but CNN may soon become the go-to tool when identification is of prime interest. Paradoxically, CNN are more computationally intensive than GMM models but may prove easier to deploy as applications and more accessible to a broad audience, as they can be trained and used directly on raw images whereas GMM approaches require meticulously prepared inputs.

Finally, if deep learning was here restricted to identification using convolutional neural networks, it has much more to offer to archaeology and morphometrics: its versatility extends to regression problems (e.g. Reese, 2021), segmentation (i.e. automating and/or improving pre-morphometrics treatment (e.g. Lee et al., 2017), adversarial reconstruction for broken or missing parts (e.g. Hermoza & Sipiran, 2018), pose and parallax correction for data acquisition (e.g. Zhang et al., 2021). In our view, this also argues for synergy rather than rivalry between CNN and GMM approaches, with future research determining the extent to which this holds true.

## Acknowledgements

This work has received funding from the European Research Council (ERC) under the European Union’s Horizon 2020 research and innovation program (Grant Agreement 852573), the French National Agency (ANR-16-CE27-0013, ANR-06-BLAN-0212-02-PHOENIX, ANR-22-CE27-0026) the International Research Program EVOLEA (France - Morocco) (CNRS-INEE) and the Défi Clé “Sciences du passé” - Occitanie region / Federal University of Toulouse, France (PATRIMOLEA programme). We are grateful to the Centre de Ressources Biologiques de la Vigne, Domaine de Vassal-Montpellier (INRAE) (https://vassal.montpellier.hub.inrae.fr) that provided all the pips from cultivated varieties and the OSU-OREME (https://oreme.org/) that helped to the constitution of the wild grape pip collection. We thank the Melgueil experimental domain (INRA, Mauguio, France), the Porquerolles (CBNMed, France), Córdoba (IFAPA, Spain) and Tessaout (INRA Marrakech, Morocco) worldwide collections of the olive tree that provide the olives stones from cultivated varieties. We also thank the Biological Resource Center-INRAE Clermont Ferrand for providing all the barley grains studied. We also want to thank Mathias Bellat, Marco Cornelli, Lloyd A. Courtenay, Thomas Huet and two anonymous reviewers for their very precious inputs during the peer-review process.

## Supplementary material

The following doi includes all scripts and data to rerun and build upon the work presented here (6.79 Go-zip): https://doi.org/10.6084/m9.figshare.25680390.v2

It includes:

* Scripts used and commented in the folder `/R`. Raw images in the folder `/DA`TA and .rda in the folder `/rda` are primarily intended to be accessed by the .R scripts.
* model histories showing accuracies, loss and learning rate for both training and validation partitions, for each combination of model x sample size x seed are in the folder `/model` histories
* hdf5 images of training weights are also available for seed 2329 in the folder `/hdf5`
* Figure 2 with the very same presentation but showing precision and recall instead of accuracy.

